# Role of livestock parameters on cattle vulnerability to drought in the coastal semi-arid areas, Kenya

**DOI:** 10.1101/2020.06.19.161273

**Authors:** Jack Owiti Omolo, Muturi Mathew, Maurice Owiny, Jeremiah N Ngugi, Joseph Ogola Ganda, Cornel K Malenga, Stephen Kioko Musimba

**Affiliations:** Zoonotic Disease Unit, Kenya; County Government of Kilifi, Department of Agriculture, Livestock Development and Fisheries; County Government of Taita Taveta, Department of Agriculture, Livestock Development and Fisheries; National Drought Management Authority (NDMA) Kilifi County; County Government of Bungoma, Department of Agriculture, Livestock and Fisheries; Kenya Field Epidemiology and Laboratory Training Program, Ministry of Health

**Keywords:** cattle, drought, vulnerability, Semi-arid

## Abstract

Livestock keeping forms main occupation in arid and semi-arid lands. Increase in drought frequency and intensity globally negatively affect livestock production and livelihood. Cattle are the most drought sensitive livestock due to size, grazing behavior and nutritional requirements. Factors for vulnerability of cattle to droughts are individual cattle parameters, health and husbandry practices. This cross sectional study aimed to those factors in semi-arid Kaloleni sub-county. Data on household (HH) head demographics, cattle and production collected from 194 enrolled HH using structured questionnaires. Cattle ages were grouped into young (<1 year old), growers (1-2 years old) and adults (>2years old). Data was analyzed using STATA 12 software. Univariate, bivariate and multivariate logistic regression analysis conducted and reported in Crude Odds Ratio (cOR), Adjusted Odds Ratio (aOR) and Confidence Interval (CI). We used Pearson product-moment correlation to determine relationship between HH head, cattle herd, individual cattle characteristics and drought characteristics, p = <0.05 being significant. Mean age HH heads was 40.7 ± 12.6 years, 44.3% (86) had basic education, males were 65.3% (n=126). Mixed livestock production was practiced by 69.1% (134), while 86.1% (167) practiced free range communal grazing. Adult cattle were 54.1% (1295). Female cattle were 72.7% (1741). Average body condition score was 3.0 ± 0.6, and calves had 2.6 ± 1.3. About 20.6% had various forms of illness, calves mostly affected at 29.1%. Up to 63.4% HH, spray cattle at home, 93.3% HH reported no vaccination history. Home straying was protective (cOR 0.3, 95% CI 0.14 – 0.53). Herd size (aOR 2.9, 95% CI 1.5 – 5.5) and having no disease control method (aOR 2.8, 95% CI 1.85 – 9.19) were contributing to reporting disease. We report positive correlation between drought outcomes and HH head (r=0.076, p>0.01), cattle herd (r=0.216, p=0.003 and individual cattle characteristics(r=0.139, p>0.01). The findings on cattle conditions exacerbate their vulnerability in presence of stressful conditions like droughts especially in calves and cows. This study demonstrates weak disease control efforts and unorganized husbandry practices. We propose strategic and focused disease control plans to improve cattle resilience and further research on livestock based factors as drought response metrics for the livestock livelihood.

## Introduction

Communities living in arid and semi-arid areas globally often face the vagaries of weather, primarily because of rise in ambient temperatures and change in rainfall patterns (4). As a result of global warming, drought has been in the last three decades reported to be increasing in frequency, duration and severity (5). The increase in frequency of drought in the arid and semi-arid lands has caused livelihood changes among agrarian communities forcing them to shift towards livestock keeping combined with crop insurance initiatives such as fast maturing and drought tolerant crop varieties production; along the seasonal river beds and valleys (6–8). In arid and semi-arid regions, livestock production is widely practiced as animals graze freely in the vast open lands under communal pastures. Livestock production has not been spared by the effects of drought associated with reduced water availability, poor quality of pasture and increase in parasite burden leading to various livestock diseases (9).

Water scarcity and challenges posed by livestock diseases in pastoral communities cause major constraints to livestock production in arid and semi-arid lands in Kenya (1).

Drought is the single most hazard which World Health Organization (WHO) estimates over 55 million people throughout the world face its wrath yearly. WHO concurs that by the year 2030, 40% of world’s population would face severe water scarcity and drought than ever before. The drought mostly affects human livelihoods in livestock and crops production (10). In sub-Sahara Africa, droughts and subsequent effects are believed to cause economic losses of up to 70% even as most African counties experience losses in Gross Domestic Product (GDP) (11). The increasing effects of drought that have led to loss of human lives, livestock deaths and crop failures have not spared countries in the horn of Africa. The region is largely arid and semi-arid having over 80% reliance on agricultural activities (12–14). A vast geographical area of Kenya is arid and semi-arid rangelands, supporting livestock production through pastoralism, leaving only 15% of land mass under crop production. Within arid and semi-arid lands, livestock production accounts for 90% of employment and household incomes (15).

Whereas livestock kept in these regions respond differently to effects of drought; cattle are the most drought sensitive livestock species based on their size, grazing behavior and nutritional requirements (8,14,16,17). Cattle are grazers making them more vulnerable to drought since pasture is a seasonal biophysical resource that is fast affected by prolonged dry weather (18). Occurrence of drought leads to increase in trekking distance, livestock migration from drier regions to places with water and pasture. Livestock migration from disease endemic areas into a new places can lead to introduction of livestock diseases to those areas and depletion of water sources and pasture (19) leading to soil erosion and conflicts.

Cattle death remain the most reported drought outcomes while other observable losses like reduced production, wasted body condition and increased disease prevalence are rarely reported (20). Vulnerability of cattle to drought outcomes also varies on individual cattle parameters and husbandry practices. Individual cattle characteristics including age, sex, breed, physiological status and production level while herd parameters like herd size and production system determine vulnerability of cattle to drought outcomes and the possibility of cattle surviving such stressful moment.

Livestock keepers do practice feed supplementation with crop leftovers, wild tree tubers and commercial feed supplements to counter the effect of drought on cattle. When drought strike, the experts often advocate for community and agency supported livestock offtake to reduce pressure for pasture and water thereby checking the loss of the livestock livelihood through deaths, abortion and other drought effects on cattle (21). The Kenyan Government has established a specialized institution in drought management, The National Drought Management Authority (NDMA), whose mandate is to establish mechanisms to end drought emergencies. These measures are established along drought cycle management which include conducting drought early warning systems, drought preparedness planning, drought contingency planning, drought response coordination and resilience building (22,23). The NDMA conducts drought early warning surveys in sentinel households within livelihood zones. The main weakness with the sentinel data collection is the lack of a structured system of collecting systematic information on livestock body condition (24). This study aimed to assess the status of parameters in individual cattle and husbandry practices, which may determine whether cattle survive in event of drought occurrence. The study attempted to estimate the cattle herd structure, body condition and production systems alongside assessing the prevailing cattle diseases reported by the disease surveillance system and control methods applied through a cross sectional prevalence study of cattle owning households.

## Methods and materials

### Study site

The study was conducted in South Eastern Kenya coast in Kaloleni Sub-county, Kilifi County (Figure 1). The sub-county has a total land mass of 686.4 Km^2^ and human population density of 227 per Km^2^ (25). Kaloleni sub-county is largely semi-arid with average temperatures recorded between 30°C and 34°C, receiving an average annual rainfall of about 300mm to 900 mm (3). The areas experience long rains during the months of March, April and May while short rains during the months of October, November and December. Rainfall is not reliable leading to poor pastures and insufficient water supply for both livestock, crop production and domestic use (22). The sub-county falls into two agro-ecological zones namely livestock-millet zone and low land ranching zone. The livestock-millet zone is mainly arid with low agricultural potential and suitable for drought tolerant crops and livestock mostly goats, sheep and cattle (27). This zone supports large numbers of livestock through free communal grazing and livestock water provision through seasonal communal earth dams and water pans which rely on rainfall for recharge (28). The low land ranching zone is mainly characterised with communal livestock grazing and commercial ranches and presence of wildlife (29). Out of seven sub-counties in Kilifi County, Kaloleni sub county holds highest numbers of livestock with over 55,000 heads of cattle, 105,000 goats, 104,000 sheep and 830 donkeys with a household owning average of 10 cattle (30).

**Figure 1.**
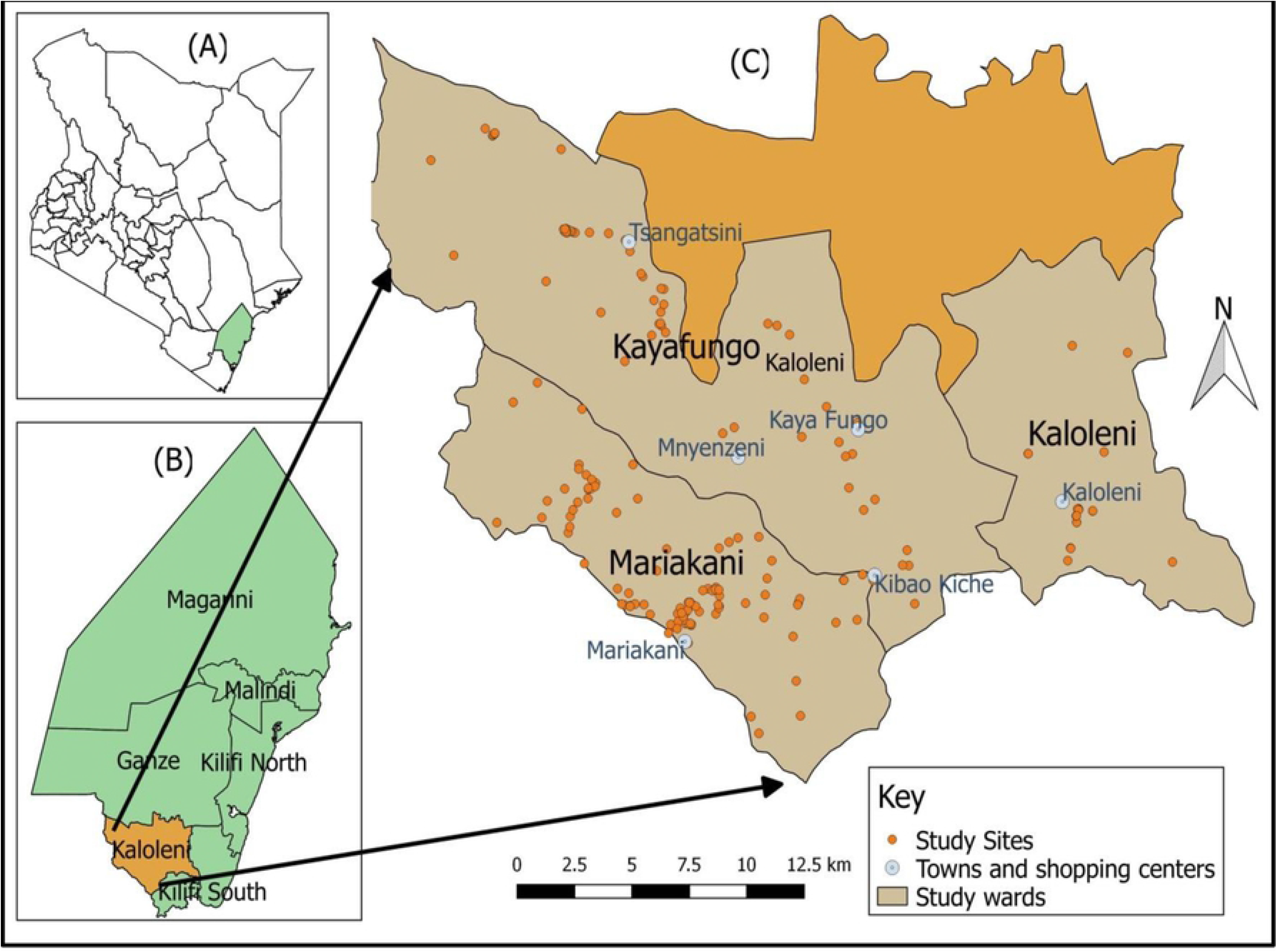
Geographical distribution of study sites, (A), map of Kenyan counties showing location of Kilifi county. (B), map of Kilifi County showing Kaloleni in relation to other sub-counties. (C), map of Kayafungo, Mariakani and Kaloleni wards showing distribution of study sites.

### Sampling strategy

This cross sectional study was conducted among the randomly selected livestock-owning households in three wards of Kaloleni Sub-county (Figure 1). A list detailing approximate numbers and distribution of households owning livestock within the three wards of Kayafungo, Kaloleni and Mariakani was obtained from sub-county veterinary and livestock officers. Stratified sampling method was used to distribute the sample size proportionate to number of households in the selected wards. In each ward, five random GPS coordinate locations were generated using QGIS software as starting points and additional two points in each ward to be used as replacements in case the initial GPS point landed in place over 1km away from nearby central point. From each starting point, the nearest central point or location was determined with assistance of local veterinary and livestock officers. The central points were defined as social centers, shopping centers, local administrators’ office or a watering point/cattle dip/vaccination crush. Sampling unit was a household with inclusion criteria of household owning cattle with or without other types of livestock. Systematic random sampling was used to identify the specific household from the household list to enroll into the study. From the identified central points within the ward, a water bottle was rotated on the ground and direction of the bottle cover was used as starting direction. The immediate household in the pointed direction was enrolled as the first household and then every fifth household was enrolled while attempting to move in straight line as possible until a maximum of seven households were enrolled in that direction. We returned to starting point and repeated the same procedure on the opposite direction to enroll households not exceeding 14 for that specific central point. The same process was conducted in other central points until sample size was obtained.

### Data collection

Sample size for enrolled households were estimated at an expected proportion (p) of livestock keeping household in semi-arid areas of 85.5% (31). To achieve this, a formula by Pfeiffer (32),

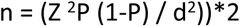

Assumptions made:
n = required sample size
Z = CL at 95% (standard value 1.96)
p = expected average prevalence of 85.5%
d = level of precision at (5%)

Minimum sample sizes of 190 households were targeted in this study. Semi-structured questionnaires were prepared and uploaded in epicollect5 software on android phones. Administration of the questionnaire was done through face-to-face interview with the household heads, who were briefed about the study and a verbal consent was obtained before administering the questionnaire. Four enumerators trained on epicollect5, principles of field surveillance, data collection techniques, did field administration of the questionnaires. Information collected included on household demographic, cattle kept, cattle herd husbandry practices. Key informant interviews with county director of veterinary services and subcounty veterinary officer was conducted together with direct observation. The GPS coordinates were collected from each sampled household using epicollect5 software.

### Applied case definitions

All the enumerators involved in this study were licensed veterinarians with ability to physically examine ill looking cattle and arrive at a tentative clinical diagnosis. A broad classification of diseases and health conditions were used for uniform conclusion by the team based on observable symptoms and clinical signs. No cattle were restrained for this purpose. Helminthes infestation was assumed to be in cattle presenting with combination of such symptoms as poor body condition, bloated abdomen, loose fecal material accompanied with history of never been dewormed in the last three months. Tick borne conditions such as anaplasmosis, babesiosis and East Coast Fever (ECF) were considered in case the cattle was anorexic, had rough hair coat, had labored breathing, had coughs or nasal discharge. foot and mouth disease and lumpy skin disease tentative diagnosis were arrived at following general examination of pathognomonic signs (33,34). Body condition was scored as described by (35) in which a scale of 0 – 5 was used with 0 being worst body condition and 5 being obese.

### Data analysis

Data from electronic devices were downloaded into MS excel spread sheet which was then uploaded into STATA 12 software for analysis. Maps were generated using QGIS-OSGeo3W 2.18.13 software. Proportions were calculated for categorical data while measures of central tendency and dispersal for continuous data. We assumed that each household enrolled owned one cattle herd. At herd level, the parameters were herd structure, cattle production and health condition. The cattle herds were grouped as large and small herds by calculating the median number of cattle from the enrolled households. Those herds with less or equal number of cattle to that of median were categorized as small herds while those with number of cattle above the median were categorized as large herds (18). The ages were grouped into young (less than one year old), growers (between one to two years old) and adults (above two years old).

Level of association between occurrence of disease in the herd as dependent variable and cattle herd size, disease prevention methods as independent variables were accessed at bivariate analysis and multivariate logistic regression. Associations were reported in Crude Odds Ratio (cOR) and Adjusted Odds Ratio (aOR) with Confidence Interval (CI) used to determine significant association. A Pearson product-moment correlation was run to determine the relationship between farmer characteristics, herd characteristics, individual cattle characteristics and drought characteristics with significant relationship having p = <0.05.

### Piloting Study

The study was piloted in Samburu ward in Kwale County, located south of Kaloleni sub-county as the two share socio-cultural environment, climatic conditions and livestock keeping trends with study area. Validity test was conducted by pilot testing, checking for concurrence of the contents and approximate time it takes to administer one questionnaire (36,37). The trained enumerators interviewed 10 livestock owners, initially about 30 minutes to administer one questionnaire. The pilot study increased brought out challenges and gaps with the study tools on reliability and consistency that were addressed and appropriate changes made.

### Ethical Considerations

Authorization was obtained from the County Director of Veterinary Services (CDVS). Verbal consent was obtained from household heads before enrollment and only those consenting were interviewed. No livestock biological samples were collected during the study. No personal identifier information of household heads was collected during this study and all data were downloaded saved in password protected computer.

## Results

### Demographic characteristics of the household heads

We enrolled 194 household heads. The mean age of household heads was 40.7 ± 12.6 years while males were 65.3% (n=126). When placed in age groups, about 30.9% (59) were young adults aged between 17 – 31 years old and 4.7% (12) were aged above 62 years. Mariakani ward had 56.0% (109) enrolled households, Kayafungo ward had 29.0% (55) and Kaloleni ward had 15.1% (29). Based on education level, 44.3% (86) of the household heads had basic primary education, 25.8% (50) had no formal education. Most 69.1% (134) of the farmers in interviewed were practicing mixed livestock farming, keeping cattle, sheep and goats in a free range 86.1% (167) (Table 1).

**Table 1.** Demographic characteristics of livestock owners enrolled for the surveillance study, the livestock kept and production system within the three wards of Kaloleni sub-county, Kilifi – 2018

| Characteristic | Location |  |  |  |
| --- | --- | --- | --- | --- |
|  | Kayafungo<br>Ward | Mariakani<br>Ward | Kaloleni<br>Ward | Total<br>n=194 |
| Household head | n (%) | n (%) | n (%) | n (%) |
| **Age groups in Years** |  |  |  |  |
| 17 – 31 | 5 (2.6) | 12 (6.3) | 42 (22.0) | 59 (30.9) |
| 32 – 46 | 12 (6.3) | 24 (12.6) | 34 (17.8) | 70 (36.7) |
| 47 – 61 | 10 (5.2) | 14 (7.5) | 28 (14.7) | 53 (27.8) |
| >62 | 0 | 5 (2.6) | 7 (2.1) | 12 (4.7) |
| **Sex** |  |  |  |  |
| Male | 36 (18.7) | 71 (36.3) | 19 (9.8) | 126 (65.3) |
| Female | 19 (9.4) | 38 (19.7) | 10 (5.2) | 67 (34.7) |
| **Level of education** |  |  |  |  |
| No formal | 1 (0.5) | 35 (18.0) | 14 (7.2) | 50 (25.8) |
| Primary | 10 (5.1) | 14 (7.2) | 62 (32.0) | 86 (44.3) |
| Secondary | 11 (5.7) | 5 (2.6) | 23 (11.9) | 39 (20.1) |
| Tertiary | 8 (4.1) | 1 (0.5) | 10 (5.2) | 19 (9.8) |

### Herd structure

From the 194 households enrolled into the study they had 2,392 head of cattle in total. The cattle were stratified into age, sex and production. Most of the cattle in the herds were adults at 54.1% (1295) followed by growers at 29.2% (699) while calves under age of one-year-old were the least at 16.6% (398). Females were the majority at 72.7% (1741). When considered into various age groups, female adult cattle were the majority in the herd at 43.5% (1041) compared to male adults, which comprised of 10.6% (254). Among the males, growers were the majority in the herds totaling to 10.8% (260) of the total cattle herds. Based on production; the cattle in Kaloleni are involved in dairy production, beef production and mixed production. Cattle keepers who practiced mixed farming were 69.1% (134), where they reared cattle for both milk and beef. The reported median milk production was at 2 liters and range of 0 – 20 liters per female. In livestock production system, free-range communal grazing method was practiced by 86.1% (167) (Table 2).

**Table 2.**
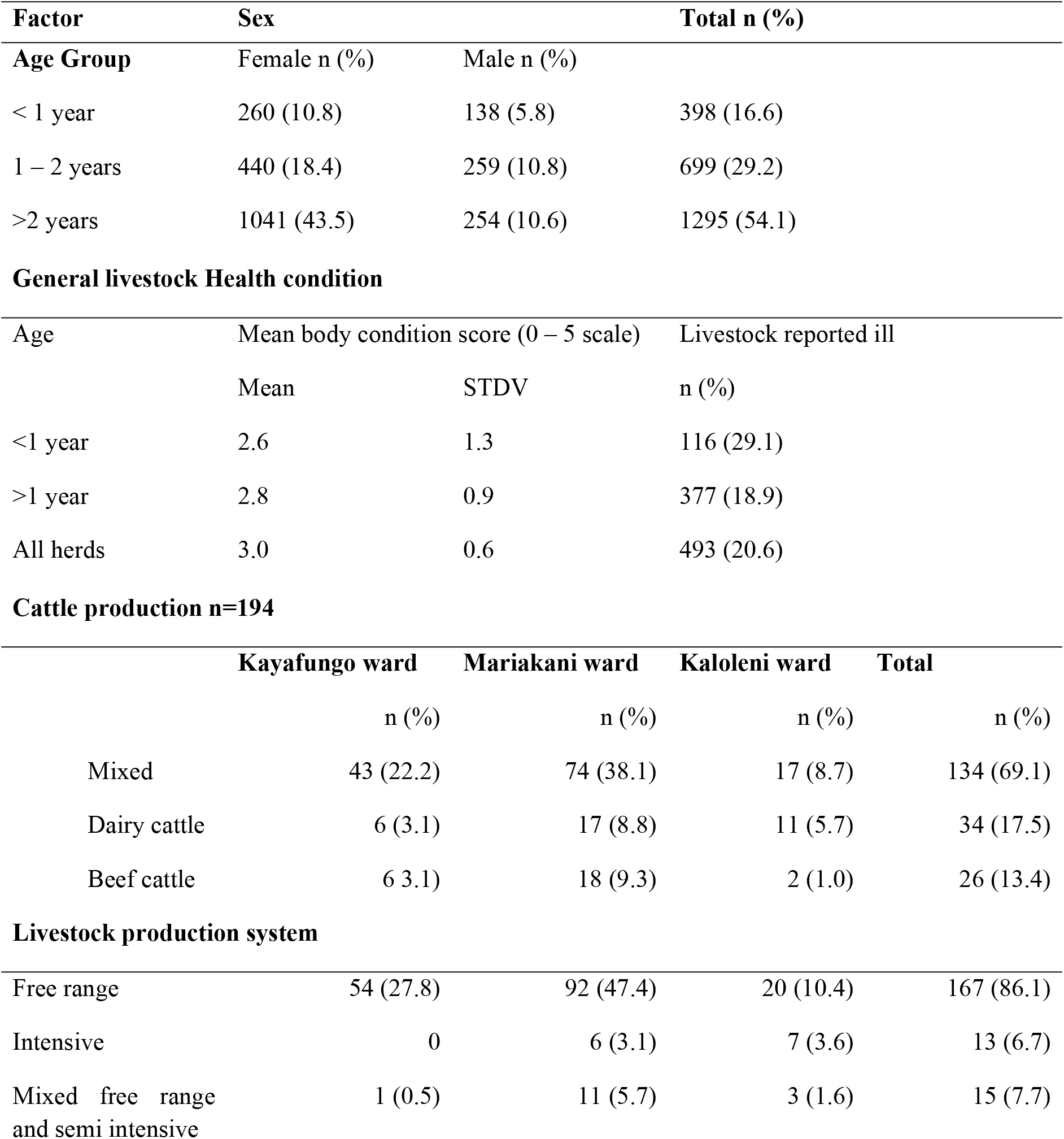
Herd structure stratified into age groups by sex, herd size and general body condition for cattle kept in Kaloleni sub-county, Kilifi County, Kenya 2018. N=2392

| Factor | Sex |  | Total n (%) |
| --- | --- | --- | --- |
| Age Group | Female n (%) | Male n (%) |  |
| < 1 year | 260 (10.8) | 138 (5.8) | 398 (16.6) |
| 1 – 2 years | 440 (18.4) | 259 (10.8) | 699 (29.2) |
| >2 years | 1041 (43.5) | 254 (10.6) | 1295 (54.1) |

General livestock Health condition
| Age | Mean body condition score (0 – 5 scale) |  | Livestock reported ill |
| --- | --- | --- | --- |
|  | Mean | STDV | n (%) |
| <1 year | 2.6 | 1.3 | 116 (29.1) |
| >1 year | 2.8 | 0.9 | 377 (18.9) |
| All herds | 3.0 | 0.6 | 493 (20.6) |

Cattle production n=194
|  | Kayafungo ward | Mariakani ward | Kaloleni ward | Total |
| --- | --- | --- | --- | --- |
|  | n (%) | n (%) | n (%) | n (%) |
| Mixed | 43 (22.2) | 74 (38.1) | 17 (8.7) | 134 (69.1) |
| Dairy cattle | 6 (3.1) | 17 (8.8) | 11 (5.7) | 34 (17.5) |
| Beef cattle | 6 (3.1) | 18 (9.3) | 2 (1.0) | 26 (13.4) |

Livestock production system
|  |  |  |  |  |
| --- | --- | --- | --- | --- |
| Free range | 54 (27.8) | 92 (47.4) | 20 (10.4) | 167 (86.1) |
| Intensive | 0 | 6 (3.1) | 7 (3.6) | 13 (6.7) |
| Mixed free range and semi intensive | 1 (0.5) | 11 (5.7) | 3 (1.6) | 15 (7.7) |

### Cattle health condition

On average, the body condition score was 3.0 ± 0.6. Calves aged less than one year had the poorest body score of 2.6 ± 1.3. About 20.6% (493) of all the cattle had various forms of illness of which calves presenting with various forms of illness were 29.1% (144). Those cattle suspected to be presenting with conditions that are tick borne were 41.0% (202) while helminthes infestation was 50.5% (259). Farmers reporting home spraying to control external parasites including ticks were 63.4% (123) while the rest took no action. Nearly 50.0% (97) of households reported seeking veterinary services whenever their cattle was sick, 25.3% (49) would treat their cattle by themselves and 24.7% (53) would call neighbor to attend to their sick cattle. Almost 93.3% (181) of the household heads had never vaccinated their herds of cattle while 64.4% (125 reported at least one cattle from their herds had died in the last three months after falling ill (Table 3).

**Table 3.** Tentative cattle diseases and conditions arrived at based on general and physical examination in Kaloleni sub-county. n=493

| **Disease/condition** | **Number observed** | **Proportion (%)** |
| --- | --- | --- |
| Total no. of cattle ill/with health conditions | 493 | 20.6 |
| Babesiosis/Anaplasmosis | 202 | 41.0 |
| Internal parasite/helminthiasis | 249 | 50.5 |
| Foot rot/Lameness | 28 | 5.6 |
| Trypanosomiasis | 14 | 2.8 |
| **Action taken by household heads when cattle is unwell n=194** |  |  |
| Action | Number | Proportion (%) |
| Call a qualified veterinarian | 97 | 50.0 |
| Treatment by household heads | 49 | 25.3 |
| Call neighbor to treat sick cattle | 48 | 24.7 |
| **Methods applied by household heads to control ticks and other vectors in cattle n=194** |  |  |
| Home spraying | 123 | 63.4 |
| Take to communal dips | 18 | 9.3 |
| No control method applied | 53 | 27.3 |
| **History of cattle vaccination against common diseases in the last 3 months n=194** |  |  |
|  | No. | Proportion (%) |
| Vaccinated | 13 | 6.7 |
| Not vaccinated | 181 | 93.3 |
| **History of occurrence of cattle disease in last 3 months leading to death of at least one cattle n=194** |  |  |
| Reported | Number | Proportion |
| Reported disease | 125 | 64.4 |
| No disease reported | 69 | 35.6 |

Various disease control methods and occurrence of disease in cattle herds were compared. Spraying at home was protective (cOR 0.3, 95% CI 0.14 – 0.53) while not practicing any disease control methods (cOR 4.2, 95% CI 2.0 – 8.85) and having large herd size (cOR 3.6, 95% CI 1.98 – 6.60) were significantly associated with occurrence of disease in cattle herd. On multivariate analysis using logistic regression, herd size (aOR 2.9, 95% CI 1.5 – 5.5) and having no disease control method (aOR 2.8, 95% CI 1.85 – 9.19) were contributing to reporting disease occurrence in cattle herds (Table 5). There was a weak positive correlation between farmer characteristics and drought outcomes, which was statistically insignificant (r=0.076, p>0.01). There was a weak, positive correlation between herd characteristics and drought outcomes, which was statistically significant (r=0.216, p=.003). There was a weak positive correlation between individual cattle characteristics and drought outcomes, which was statistically insignificant (r=0.139, p>.01) as shown in (Table 6).

**Table 4.** Annual report from sub-county veterinary office on cattle diseases and control measures in Kaloleni, Kilifi County – 2017/2018

| Disease Reported |  | Number of cases attended |
| --- | --- | --- |
| ECF |  | 920 |
| Anaplasmosis |  | 416 |
| Babesiosis |  | 62 |
| Trypanosomiasis |  | 326 |
| Helminthes |  | 1615 |
| Pneumonia |  | 373 |
| Cattle vaccinations |  | Number vaccinated |
| RVF |  | 3678 |
| Anthrax and black-quarter |  | 6224 |
| Tick control measures | No. of operational units | No of cattle reached |
| Communal dips | 1 | 1123 |
| Groups doing home spraying | 7 | 2480 |

**Table 5.**
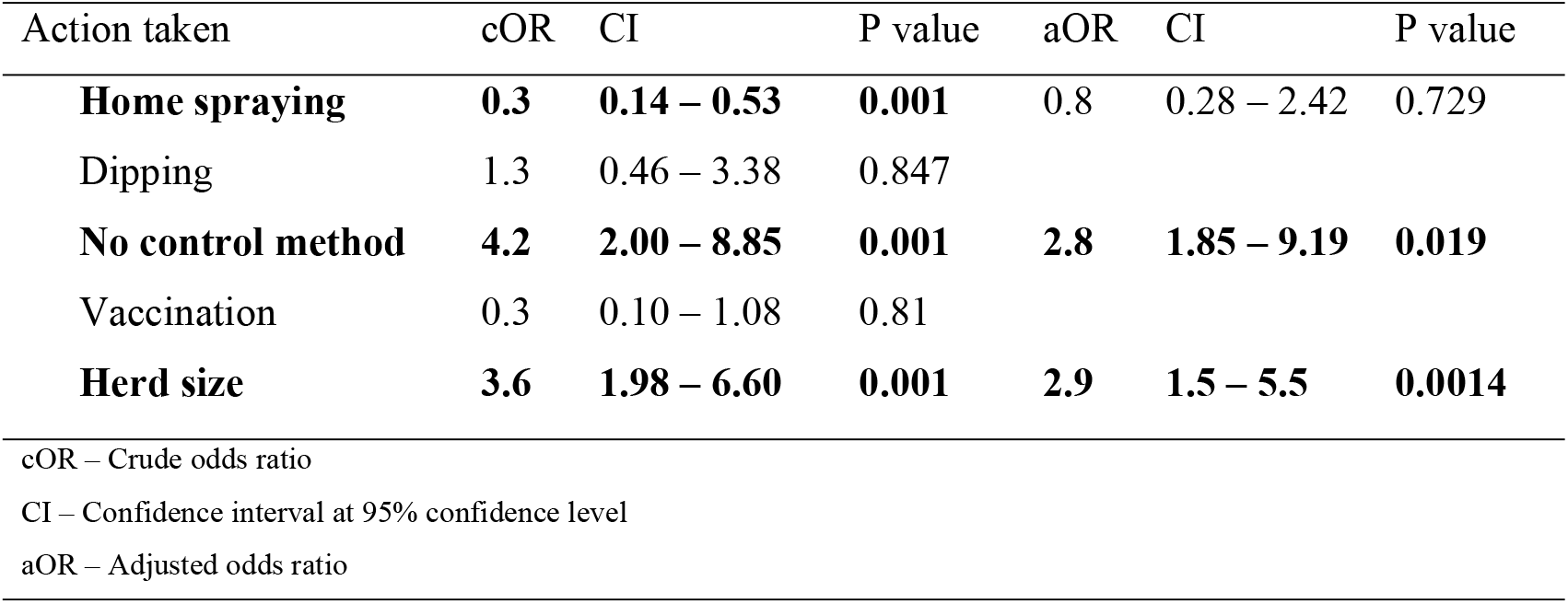
Association of cattle disease prevention methods and herd size to occurrence of disease in cattle herd in the past three months in Kaloleni sub-county, 2018

**Table 6.** Multivariate analysis on correlation between farmer characteristics, herd characteristics, individual cattle characteristics and how they are influenced by drought characteristics in the past three months in Kaloleni sub-county 2018. (n=194)

|  |  | Household<br>head<br>Characteristics | Cattle Herd<br>Characteristic | Individual<br>Cattle<br>Characteristic | Drought<br>Characteristics |
| --- | --- | --- | --- | --- | --- |
| Farmer Characteristics | Correlation Coefficient | 1.000 | **\*\*0.202** | **\*\*0.201** | 0.076 |
|  | P Value | . | **0.005** | **0.005** | 0.290 |
| Herd Characteristic | Correlation Coefficient | **\*\*0.202** | 1.000 | **\*\*0.956** | **\*\*0.216** |
|  | P Value | **0.005** | . | **0.000** | **0.003** |
| Individual Cattle Characteristic | Correlation Coefficient | **\*\*0.201** | **\*\*0.956** | 1.000 | 0.139 |
|  | P Value | **0.005** | **0.000** | . | 0.053 |
| Drought Characteristics Characteristic | Correlation Coefficient | 0.076 | **\*\*0.216** | 0.139 | 1.000 |
|  | P Value | 0.290 | **0.003** | 0.053 | . |

**\*\*** Significant correlation

### Key informant interview

Reports at the sub-county veterinary offices show that most prevalent cattle diseases in the area are trypanosomiasis and tick borne diseases (ECF, babesiosis and anaplasmosis) of which the government prevention and control mechanisms reported were communal dipping, routine passive disease surveillance and ambulatory response to sick cattle. Vaccine preventable diseases reported were Foot and Mouth Disease (FMD), Lumpy Skin Disease (LSD), Rift Valley Fever (RVF) and anthrax. In the previous year prior to this study, subcounty veterinary office conducted disease prevention through vaccination against RVF and Anthrax. This was through public vaccination campaigns funded by both county government and development partners. The tick and parasite control methods involved dipping and provision of foot pumps and acaricide to farmers. Of the seven rehabilitated cattle dips, only one was operational as lack of reliable water provision and local management mentioned as main challenges hindering dip operations. On direct observation, pasture condition was fair to good with those in public land communal grazing fields being of low quality and quantity that ones in enclosed privately owned lands. Several herds of cattle were observed grazing together in a shared grazing field. Water source was adequately reliable at the time of study, comprising of seasonal rivers and earth dams, which rely heavily on rainfall and are approximated to last three months if not recharged (Table4).

## Discussion

### Demographic characteristics of household heads

Most of those interviewed were youthful males possibly because males are usually the heads of households and decision makers in African settlements. Despite females being mostly involved in agricultural activities globally and in Africa, men are usually seen as livestock asset owners and decision makers (38,39). Most of those interviewed had completed primary level of education and another quarter reporting no formal education. This finding contradicts those from other studies which indicate majority of livestock keepers in arid and semi-arid lands lacks any form of education (40). Such findings apply to nomadic pastoralists inhabiting large underdeveloped areas with no infrastructure network as opposed to Kaloleni sub-county in which a major road highway that form part of the northern corridor from Mombasa city to Nairobi city exits. The inhabitants do not practice any form of pastoralism hence social amenities including those of education are available (25). Level of education affects the capability of an individual in making critical decisions particularly in times of drought and adapting to new farming technologies (41). Arriving at a decision during stressful and complex situations as severe drought, an individual not only rely on level of education but also known traditional practices, exposure levels to other options and nearby practices (41) hence the routine extension and continuous education may be useful in decision making during stress. Gender and level of education are among such factors that have been identified as significant in influencing the response to drought by the cattle keepers (42).

### Cattle parameters and husbandry

All cattle herds were stratified into age, sex and production systems. Majority of the cattle in all the herds were female most of which were above two years old. These females form the significant economic part of production as they provide milk for both farm subsistence use, feeding their young ones and for sale. Females often stay longer in the herds than males and are disposed normally due to reduced production, aged or as a result of chronic disease (15). This longer stay of females creates sentimental and functional relationships with the holders. Their longevity in the herd expose them to drought outcomes such as migratory diseases (43) and have to endure several episodes of pasture and water shortages. Majority of the males were of mid age between 1 – 2 years old while those under one year were few. The males were either born within the herd or introduced into the herd for fattening reasons. Studies have shown that farmers do purchase males from livestock markets for fattening during wet seasons when pasture and water are available (44). The cattle herd structure results were similar to the one in pastoral arid regions in north eastern Kenya (45), however, the herd volumes were larger in pastoral region than that observed coastal semi-arid region. This observation was attributed to vast communal land in northern Kenya with pastoralists whose main occupation is livestock keeping. In coastal region, low cattle herd volumes are due to limited land, cultural practices where communities have settled in specific places and availability of other sources of household income (25).

Body condition of cattle predicts how well it can survive through a nutritional scarcity moment (1). The body condition is direct indicator of the health of cattle and is dependent on individual cattle physiological status and cattle husbandry practices (37). The general body condition in the herds enrolled into this study ranged from average to good. During this time, the herds were recovering from a dry period. Calves had lowest body condition score when compared to older cattle. The young cattle in the herd mostly graze together with adult ones. This exposes them to competition with adults for the available pasture. The calves which are yet to be weaned rely on milk from their dams which is shared with livestock keepers (46).

These factors can be responsible for the low body condition in under one-year-old animals. In an event of drought, the calves are likely to suffer most in the face of limited water and pasture. Cattle production is another aspect which has been reported to be severely affected by drought occurrence (23). In Kaloleni sub-county, most of cattle keepers practice mixed cattle production where keeping of dual-purpose zebu cattle for meat, milk and cattle sales. Such cattle breeds are preferred by local farmers because they are believed to be more tolerant to drought outcomes despite low production and growth rates (18). Most cattle keepers were practicing free-range communal grazing methods on community and public owned lands and along the road reserves. The water sources were mainly earth dams and seasonal rivers that rely on rainfall reliability and intensity. This husbandry practices has been associated with over grazing, transmission of livestock diseases and lack of grazing control and intercommunity conflicts due to scarcity of resources (3). In event of drought occurrence, such livestock production system leads to acute depletion of pasture and water thus exposes the cattle to severe effects of drought. Use of communal land for grazing has been associated with severe cattle losses in event of drought occurrence as it inflates the already existing pressure pasture and water (47).

We observed that up to quarter of cattle in herds had signs of disease or poor health conditions. Most of these observations were among the calves and female. Young cattle have not fully developed immune system to fight local prevailing livestock diseases. The young cattle require special nutritional attention for healthy growth (48). In a possibility of drought occurrence, the young cattle less than one year old could be worst affected (49). Drought leads to reduced pasture quality and quantity, thus the individual cattle fail to meet the nutritional requirements. This weaken intrinsic body protection mechanisms against prevailing diseases in the face of increasing parasites and disease incidences (50). The mitigation mechanisms of drought involving livestock migrations and mixing up of herds which share communal grazing and watering points increase the incidences of livestock pests and diseases (19,48,51). Babesiosis, anaplasmosis and East Coast Fever (ECF) diseases form majority diseases tentatively diagnosed from cattle herds in the study. A study in Zimbabwe by Tukanda 2019 (40), observable increase in livestock diseases ranked high as a consequence of drought in cattle. This may be due to opportunistic nature of infectious agents, increased interaction between the herds in communal watering and grazing fields. Strategically planned livestock disease control measures by both local veterinary authorities and livestock farmers should have drought cycle in mind. Based on our study findings from households, high proportion of cattle keepers in Kaloleni sub-county had never had their cattle vaccinated against any disease in the previous year. This might be responsible for the reports of disease preventable cattle disease leading to morbidity and deaths. Successes of such program rely on coordination and adequate information sharing between livestock keepers and implementers of such program (52). Successful livestock vaccination program requires high coverage targeting over 70% of susceptible livestock (53). Properly implemented livestock vaccinations have been demonstrated to offer economic advantages as disease occurrences reduce hence more healthy livestock and human population (52,54,55). During drought incidences, government and its development partners often conduct mass livestock vaccination as a measure of mitigation. This has been found to have no effect on deaths and reports of livestock diseases (56) while sustained livestock vaccination is beneficial (57).

The numbers of cattle keepers using communal dipping were minimal. This was attributed to low number of rehabilitated and active cattle dips within the sub-county, the distance between the household to the cattle dip, and the amount of levy paid to dip management committee. A study conducted in Tharaka Nithi County in central Kenya found that community sensitization and properly trained cattle dip management improved performance of cattle dips and proposed initial provision of acaricide and water by the government followed by slow handing over process (58). Payment of fees to access veterinary services have been shown as hindrance to livestock keepers from benefiting from such services (8,29). As a coping mechanism, majority of the cattle keepers practiced home spraying as a way of controlling vectors and other external parasites. This method has been found to be most preferable for its convenience and considered cheap (19). It however comes with challenges of getting right dosage, application skills, safety of the user and environment. It may lead to overuse of a particular acaricide resulting in possible tick and other parasite resistance (60). Various methods of applying acaricides to cattle present different challenges and success with more efficient ones being more expensive and out of reach to most cattle keepers in arid and semi-arid areas. Spraying at home, however suitable it could be to most cattle keepers in Kaloleni sub-county, has environmental and efficiency concerns (61). A quarter of cattle keepers in Kaloleni sub-county do not practice any form of external parasite control. Likelihood of cattle disease occurrence in a herd was significantly associated with herd size. Cattle raised in a large herd were likely to report disease as chances of one being ill and transmitting to others. We significantly observed correlation between drought outcomes to that of household head; cattle herd and that of individual cattle.

### Conclusion and recommendation

The existing conditions at individual cattle, herd level and husbandry practices exacerbate the cattle vulnerability in presence of stressful conditions like droughts. Disease control and prevention methods applied seem to be weak and do not provide adequate protection of cattle against prevailing diseases as diseases and livestock parasites are endemic in cattle herds in the semi-arid areas. Strategic and focused disease control plans to reduce morbidity and mortality is necessary to improve cattle resilience. Availability of relatively fair road network and nearby developed towns is opportunities that can be exploited to improve the cattle health and welfare hence improving household income and livelihood.

The study recommends sustained, subsidized livestock extension services, targeting good agricultural production practices by the county government and relevant partners, on actions that can improve health and body condition of cattle like feed conservation and disease prevention and control. The findings form important source reliable information to both local and national governments for formulating activities and policies towards improving cattle production, veterinary services and increasing drought resilience. In order to enhance the sensitivity of drought monitoring, NDMA should standardize and incorporate livestock parameters into drought early warning system. The study also recommends further research on livestock based factors as drought response metrics for the livestock livelihood.

## Acknowledgement

We appreciate and value the support we got from the County Director of Veterinary Services (CDVS) of Kilifi County for allowing his officers to participate in the implementation of the study. Directorate of veterinary services at Kaloleni Sub-county was of great help in the field and we are sincerely grateful to them. We also acknowledge NDMA Kilifi County for their technical support in developing and implementing the study. This study could not have been complete without full support from local administrators and household communities in Kaloleni sub-county, whom we are greatly indebted.

## Author contributions

JOO, SKM, Participated in conception of the study idea, proposal writing, implementation, analysis and writing and review of manuscript

CKM, Conception of study idea and review of manuscript

MM, MO, JNN and JOG, Review of manuscript

